# Macrophages are required to coordinate mouse digit tip regeneration

**DOI:** 10.1101/104356

**Authors:** Jennifer Simkin, Mimi C. Sammarco, Luis Marrero, Lindsay A. Dawson, Mingquan Yan, Catherine Tucker, Alex Cammack, Ken Muneoka

## Abstract

In mammals, macrophages are known to play a major role in tissue regeneration. These cells contribute to inflammation, histolysis, re-epithelialization, re-vascularization and cell proliferation. While macrophages have been shown to be essential for epimorphic regeneration in salamanders and fish, their role has not been elucidated in mammalian epimorphic regeneration. Here, using the mouse digit tip model as a mammalian model of epimorphic regeneration, we demonstrate that macrophages are essential for the regeneration process. Using cell depletion strategies, we show that regeneration is completely inhibited; bone histolysis does not occur, wound re-epithelization is inhibited and the blastema does not form. Rescue of epidermal wound closure, in the absence of macrophages, promotes blastema accumulation but not differentiation indicating that macrophage play a role in re-differentiation. Additionally, inhibition of osteoclasts and the degradation process is not sufficient to inhibit regeneration. These findings show that macrophages play an essential role in coordinating the epimorphic regenerative response in mammals.

## INTRODUCTION

While all animals display some level of regenerative ability, some just do it better. Among vertebrates, urodele amphibians possess the ability to faithfully regenerate large parts of their body, for example their limbs (Brockes, 1997), while a number of fish species including zebrafish, readily regenerate their tail fins (Gemberling et al., 2013; Pfefferli and Jazwinska, 2015). These examples involve the coordinated regeneration of multiple tissues. This process is mediated by the formation of a blastema, a structure of proliferating undifferentiated cells that re-utilize developmental mechanisms to replace lost structures (Brockes and Kumar, 2002; Bryant et al., 2002; Tanaka, 2003). Blastema-mediated regeneration is called epimorphic and is considered to be distinct from the regeneration of individual damaged tissues, such skin and bone, which undergo a repair response without forming a blastema (Carlson, 2005). Characteristics of epimorphic regeneration, mostly gleaned from studies of model systems such as the urodele limb or the zebrafish fin, include rapid re-epithelization, formation of a signaling wound epidermis, requirement for neurotrophic factors, and the involvement of a heterogeneous lineage-restricted population of blastema cells that can re-enter the cell cycle (Stocum and Cameron, 2011).

Among mammals there are only a few models of epimorphic regeneration and the mouse digit tip is the best characterized (Borgens, 1982; Fernando et al., 2011; Han et al., 2008; Neufeld and Zhao, 1993). Digit tip regeneration in mice parallels the regeneration of human fingertips, a process well documented in the clinical literature (Illingworth, 1974; McKim, 1932), and displays some characteristics that are similar to amphibian models of limb regeneration (Han et al., 2005; Muller et al., 1999; Muneoka et al., 2008). The mouse digit tip consists of the third phalangeal element, surrounded by loose connective tissue and encased within the nail organ (Simkin et al., 2015a). The regeneration response can be divided into three distinct but overlapping phases: wound healing, blastema formation and re-differentiation. Wound healing is characterized by an osteoclast-driven bone degradation response that precedes wound closure, thus re-epithelization of the amputation wound is delayed by comparison to other epimorphic regeneration models (Fernando et al., 2011; Han et al., 2008). The completion of wound closure signals the termination of bone degradation and the transition to the blastema formation phase (Simkin et al., 2015b). Cell lineage studies provide evidence that the blastema is composed of a heterogenous population of lineage-restricted cells (Lehoczky et al., 2011; Rinkevich et al., 2011), which are actively recruited to the amputation site (Lee et al., 2013). The blastema is avascular and hypoxic; two characteristics essential for a normal regenerative response (Sammarco et al., 2015; Sammarco et al., 2014; Yu et al., 2014). Re-differentiation of the blastema to regenerate the amputated P3 bone occurs by direct ossification and is dependent on BMP and WNT signaling (Lehoczky and Tabin, 2015; Takeo et al., 2013; Yu et al., 2010). Amputation of digits at more proximal levels fail to regenerate but results in a periosteal response that forms a bone callus similar to a fracture healing response (Dawson et al., 2016).

The primary regenerated tissues during epimorphic regeneration of digit tips are skin and bone. The repair of mammalian skin has been studied primarily in full thickness wounds in which the healing response consists of distinct but overlapping phases beginning with hemostasis/inflammation, the formation of granulation tissue and finally, matrix remodeling (Eming et al., 2014). The inflammatory response dominates the early stages of healing and is critical for re-epithelization as well as supporting granulation tissue formation (DiPietro et al., 1998; Goren et al., 2009; Leibovich and Ross, 1975; Mirza et al., 2009). During matrix remodeling, the immature scar tissue deposited by granulation tissue is realigned and cross-linked to form the mature scar (Xue and Jackson, 2015). In general, skin wound healing is considered non-regenerative, and because inhibiting macrophages results in scar-free healing, inflammation plays an inhibitory role in skin regeneration (Ashcroft et al., 1999; Martin et al., 2003; Mori et al., 2002). Alternatively, tissue-specific bone regeneration has been studied in the context of fracture healing and consists of distinct overlapping phases that begins with inflammation and ends with remodeling of the regenerated bone (Schindeler et al., 2008). Macrophage depletion studies show that these cells are required for bone regeneration and successful fracture healing (Alexander et al., 2011; Raggatt et al., 2014). Thus, in this regeneration model inflammation plays an essential and stimulatory role in the regeneration of new bone.

It is not intuitively obvious how the epimorphic regenerative properties of the digit tip relate to the tissue specific repair properties of bone (regenerative) and skin (non-regenerative). Nevertheless, both repair responses initiate with inflammation; in the case of wound healing, its action is inhibitory whereas in fracture healing its action is stimulatory. Inflammation in mammalian epimorphic regeneration has not been investigated although recent studies in urodele limb and zebrafish fin regeneration show that macrophages are required for regeneration (Godwin et al., 2013; Petrie et al., 2014). In this study we examine the inflammatory response of epimorphic regeneration in mammals. Enhancing macrophage numbers, in the amputated mouse digit tip, does not have a major effect on the regeneration response; however depleting macrophages during the early stages of regeneration inhibits regeneration. Macrophage depletion inhibits osteoclast recruitment, bone degradation, wound re-epithelization, and blastema formation. Specific depletion of osteoclasts coupled with rescuing wound re-epithelization inhibits bone degradation, but a blastema forms and regeneration ensues. Thus, while osteoclast-driven bone degradation is dependent on macrophages, this process is not required for the regeneration response. Coupling macrophage depletion with the rescue of wound re-epithelization stimulates the formation of a distal blastema-like structure but does not rescue regeneration. With these data we conclude that formation of the wound epidermis is a macrophage-dependent process required for the distal recruitment of cells to form a blastema. Finally, independent of a role in osteoclastogenesis or wound re-epithelization, macrophages are necessary for the re-differentiation stage of regeneration. These results show that macrophages play critical roles in coordinating activities during all three phases of epimorphic regeneration in mammals.

## RESULTS

### Both neutrophils and macrophages accumulate at the injury site after digit amputation

Amputation of an adult mouse terminal phalangeal element (P3) transects the nail plate, epidermis, dermis including blood vessels and nerves, and bone (Simkin et al., 2013). Like all traumatic injuries, P3 amputation initiates an inflammatory response that involves the influx of cells derived from the hematopoietic cell lineage and, since these cells are not present in regenerated tissues (Rinkevich et al., 2011), they represent a transient cell population. To characterize this inflammation response, immunohistochemical techniques were used to analyze the timing and position of CD45^+^ hematopoietic cells, Ly6B.2^+^ neutrophils (Hirsch and Gordon, 1983) and F4/80^+^ macrophages (Austyn and Gordon, 1981) within the amputation wound (Fig. 1A). Prior to and immediately following amputation, there are few neutrophils or macrophages present in the mature digit (Fig. 1B,F). Indeed, there are few CD45 positive cells in the mature digit indicating that the pool of resident cells of the hematopoietic cell lineage prior to amputation injury is very low (Fig. 1J). Following amputation we find a progressive influx of neutrophils within digit stump tissues that peaks at 5 days post amputation (DPA) (ANOVA main effect time, F= 10.54 p=0.0002; *Bonferroni post hoc test p<0.05). Ly6B.2^+^ cells associated with the scab at 3 DPA appear to be dead or dying, while neutrophils present in the stump are mainly present in the bone marrow cavity (Fig. 1C arrows). At 7 DPA the wound epidermis is not yet closed; neutrophils are localized to the bone marrow and the dermal connective tissue surrounding the bone stump (Fig 1D). When the blastema forms by 10 DPA, neutrophils are predominately localized to the blastema and the bone marrow cavity (Fig 1E). By 15 DPA, neutrophil numbers return to pre-amputation levels (Fig. 1A).

**Figure 1.**
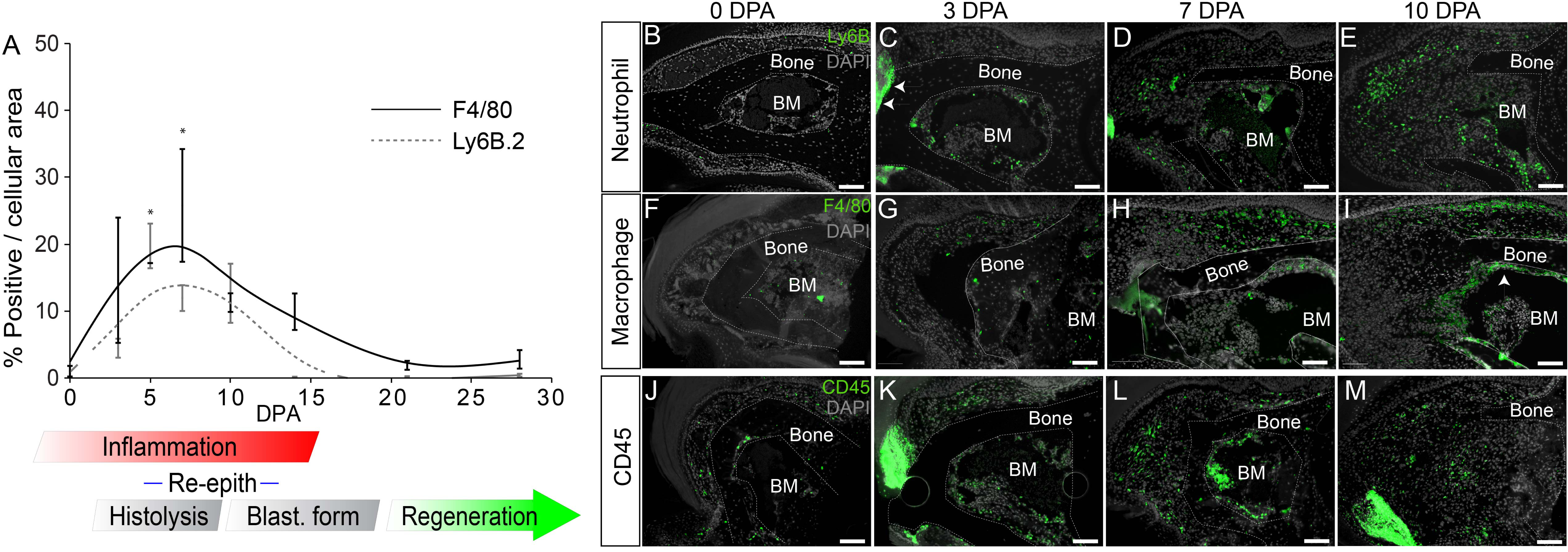
Leukocytes are recruited to the injury site after amputation. **(A)** Macrophage and neutrophils numbers are quantified using immunohistochemistry for the pan-macrophage marker, F4/80, and the neutrophil cell surface protein, Ly6B.2. Both macrophage and neutrophil numbers increase at the wound site temporarily following a regenerative P3 amputation. Stages of regeneration are delineated beneath the graph: Inflammation, Histolysis, Re-epithelization, Blastema formation, and Regeneration. (*Bonferrori post hoc test p<0.05 for Ly6B.2 0 DPA vs. 5 DPA and F4/80 0 DPA vs. 7 DPA. n=3/timepoint). **(B-E)** Ly6B.2 + cells (green) are observed in the injury area after amputation, in low numbers at 0 DPA (B), in the bone marrow and in scab (white arrows) at 3 DPA (C), and in the forming blastema and bone marrow at 7 DPA (D) and 10 DPA (E). **(F-I)** F4/80+ macrophages (green) are observed in low numbers at 0 DPA (F) and in the soft connective tissue surrounding the bone and bone marrow at 3 DPA (G) and 7 DPA (H), with bone marrow lining macrophages becoming more evident at 10 DPA (I, white arrowhead). **(J-M)** Presence of CD45 + leukocytes (green) surrounding the P3 digit is low at 0 DPA (J) and increases after amputation at 3 DPA (K), 7 DPA (L) and 10 DPA (M). Grey = DAPI nuclear stain, Scale bars = 100μm. DPA = days post amputation. BM = Bone marrow. For all images: distal = left, dorsal = top

Following digit amputation, macrophage numbers peak at 7 DPA and return to baseline by 21 DPA (Fig. 1A, ANOVA with main effect time, F=3.18, p=0.04, *Bonferroni post hoc test p<0.05). F4/80^+^ cells are seen in low numbers in the bone marrow immediately following amputation (Fig 1F). At 3 DPA, macrophages are scattered within the bone marrow and also in the connective tissue surrounding the digit stump (Fig 1 G). At 7 DPA, macrophages are predominately found within the dermis associated with the nail matrix in the proximal portion of P3. By 10 DPA, macrophages associated with the nail matrix remain high and at this stage, F4/80^+^ cells line the endosteal layer of the P3 bone marrow (Fig. 1H,I). Notably, few F4/80^+^ macrophages are observed in the blastema itself. Immunohistochemical staining of the pan-hematopoietic marker CD45 at similar regeneration stages identifies cells within the bone marrow, stump dermis, blastema that overlap the combined staining for neutrophils and macrophages (Fig. 1K-M) and suggests that these cell types represent the majority of the hematopoietic response to digit amputation. Overall, the regeneration response is associated with an accumulation of neutrophils found predominately within the bone marrow and blastema, while macrophages are localized to the endosteum of the P3 bone and the stump dermis associated with the nail matrix.

### Macrophages are required for digit tip regeneration

To explore the role that the macrophage population has in digit tip regeneration, we first tested the hypothesis that increasing macrophage presence inhibited regenerative capacity. We used targeted application of Macrophage Chemoattractant Protein (MCP-1/CCL2) following digit tip amputation to enhance the recruitment of activated macrophages (Dipietro et al., 2001). A micro-carrier bead soaked in a high concentration of MCP-1 (0.5µg/µl) was implanted in the connective tissue of the P3 digit. In uninjured digits this MCP-1 treatment is able to enhance macrophage recruitment over control BSA treated beads at 5 days post implantation (DPI), and macrophage levels returned to control levels by 15 days (Fig. 2A, *Bonferroni post hoc test, main effect treatment, p<0.05). When MCP-1 treatment was coupled with digit amputation, we observe a higher influx of macrophage numbers when compared to BSA-treated digits at 5 and 15 DPA (Fig. 2A, *Bonferroni post hoc test, main effect treatment p<0.05). The enhanced macrophage presence was observed in both the dermis and bone marrow (Fig. 2B, C). These data show that MCP-1 treatment successfully enhances and sustains macrophage recruitment to the regenerating amputation wound. Following the regeneration process using μCT *in vivo* imaging, we found that MCP-1 treated digits successfully regenerated largely in parallel with controls (Fig. 2D, E). Bone volume measurements during the regenerative response indicated a statistically significant reduction in bone degradation associated with MCP-1 treatment (Fig 2D *Bonferroni post hoc, main effect treatment p<0.05). This result was not anticipated based on evidence that MCP-1 enhances osteoclast differentiation in vitro and also enhances foreign body giant cell fusion *in vivo* (Khan et al., 2016; Kyriakides et al., 2004). Nevertheless, the findings indicate that enhancing macrophage recruitment at the digit amputation wound does not inhibit the regenerative response. These data do not support the hypothesis that macrophages are inhibitory for regeneration in mammals.

**Figure 2.**
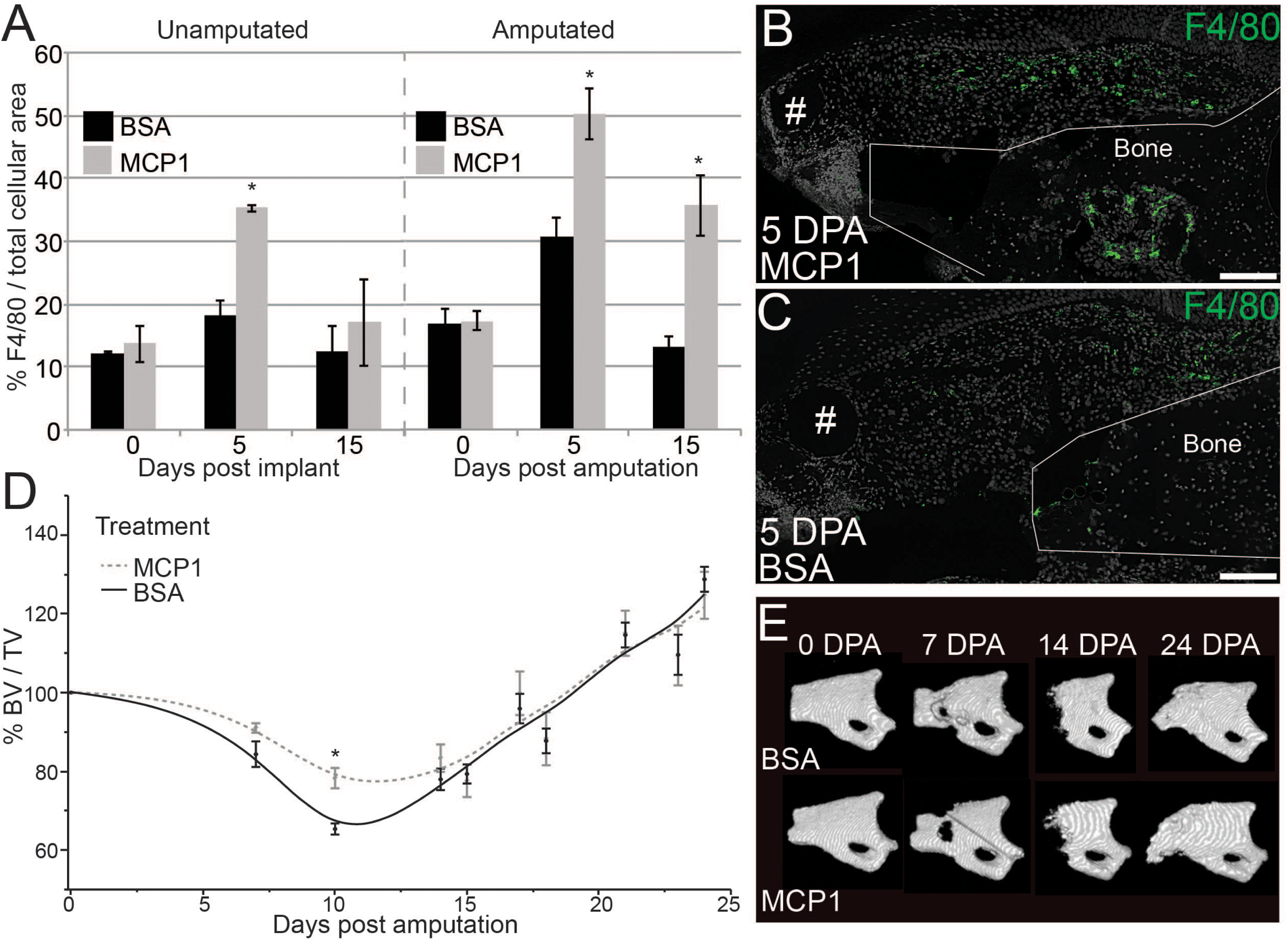
Increasing macrophage numbers does not inhibit regenerative ability. **(A)** The introduction of monocyte chemoattractant protein-1 (MCP1) via a microcarrier bead is able to recruit a higher number of F4/80+ cells to P3 when compared to BSA control beads both in unamputated digits and following amputation. Y-axis = total area of F4/80 + signal per total area of DAPI + signal in the connective tissue area. X-axis = days post bead implantation in unamputated and amputated digits (n=7 mice, 14 digits per treatment, *Bonferrori post hoc test p<0.05 for main effect treatment) **(B-C)** Immunofluorescence with anti F4/80 at 5 DPA after implantation of microcarrier beads soaked in MCP1 (B) or BSA (C). Green= F4/80, Grey = DAPI, # = microcarrier bead. Scale bars = 100μm **(D)** Micro-CT analysis of bone volume change over time. MCP1 treated digits show a significant reduction in amount of bone degradation when compared to BSA controls, but show the same overall bone volume growth by DPA 24 as compared to BSA controls. Y-axis = percent bone volume per total volume at time of amputation (%BV/TV). X-axis = Days post amputation. (n=7 mice, 14 digits per treatment, *Bonferroni post hoc test p<0.05 for main effect treatment). **(E)** 3D renderings of μCT data show patterned bone growth in both BSA and MCP1 treated digits by 24 DPA. For all images: distal = left, dorsal = top

To explore the effect of depleting macrophage numbers during digit tip regeneration we used a commercially available reagent, Clodronate Liposomes, that is effective in transiently depleting macrophages when applied either systemically or locally (Alexander et al., 2011; Barrera et al., 2000; Li et al., 2013; Xiang et al., 2012). Clodronate Liposomes are selectively engulfed by phagocytic cells and are cell lethal; however unengulfed liposomes are rapidly cleared via the kidneys (van Rooijen and Hendrikx, 2010). We used a local Clodronate Liposome treatment of individual mouse digits during the rise in macrophage recruitment to test whether phagocytic cells are required for digit tip regeneration. Clodronate Liposomes (Clo-Lipo) or control liposomes containing PBS (PBS-Lipo) were injected into amputated digits at 0, 2 and 5 DPA. Samples were collected at 6 DPA to check for neutrophil and macrophage presence based on immunohistochemistry and for osteoclast presence based on distinct cytological characteristics, i.e. multinucleated giant cells with ruffled borders. We observed a significant reduction in F4/80^+^ cells in Clo-Lipo injected digits as compared to PBS-Lipo controls indicating that targeted treatment effectively diminished the local macrophage population (Fig. 3A, *unpaired student's t-test p<0.05). Osteoclasts which are derived from the monocyte/macrophage lineage and are known to be present in high numbers during regeneration, (Fernando et al., 2011; Sammarco et al., 2014), were also depleted while neutrophils and osteoblasts are largely unaffected by Clo-Lipo treatment (Suppl. Fig. 1). These findings show that Clo-Lipo treatment is effective in locally depleting the injury-induced macrophage population.

**Figure 3.**
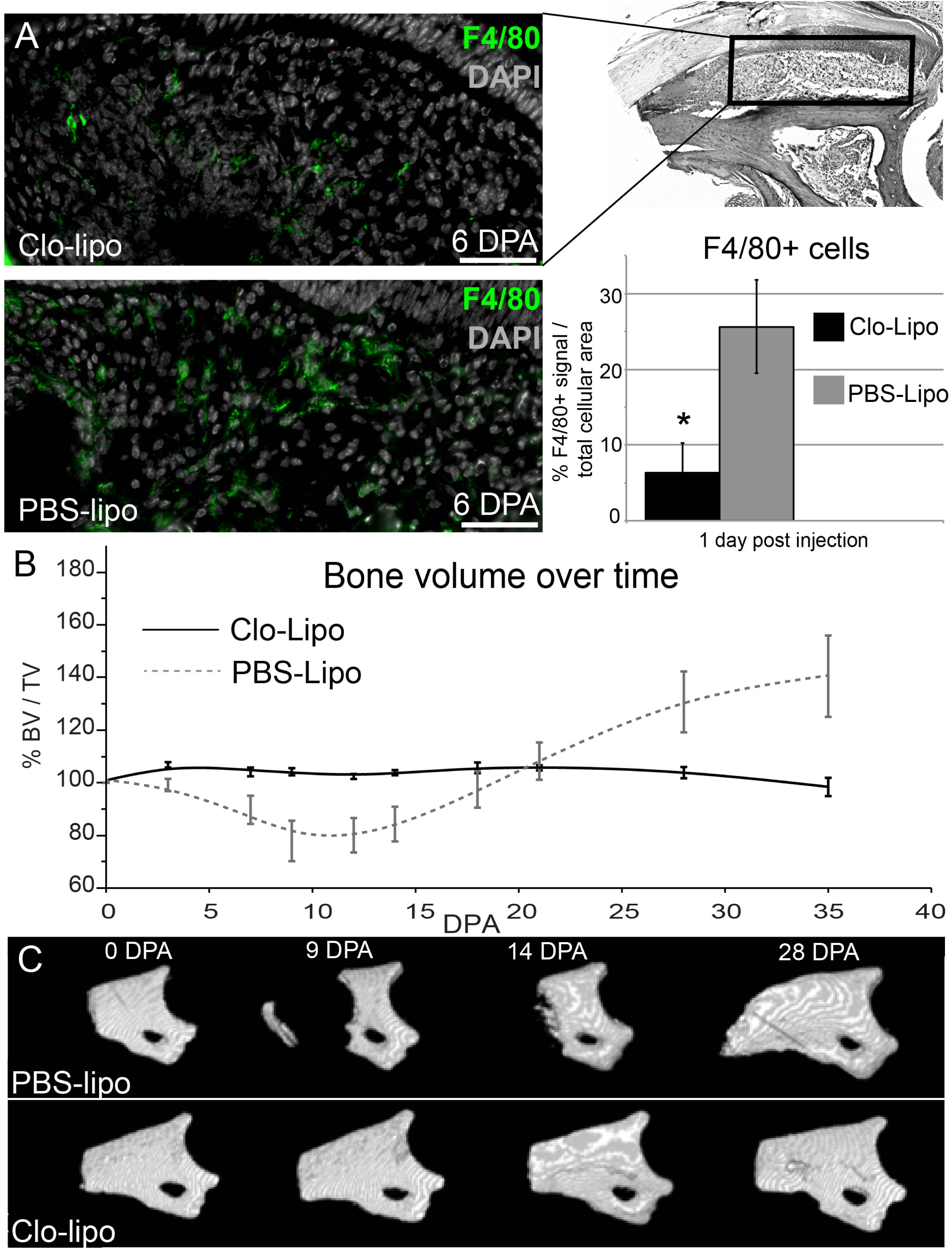
Local injections of clodronate liposomes effectively deplete macrophage populations and inhibit regeneration. **(A)** 50μg of either clodronate liposomes (Clo-Lipo) or PBS liposomes (PBS-Lipo) were locally injected into the P3 digit at days 0, 3, and 5 after amputation. Quantification of F4/80+ cells at 6 days post amputation (1 day post final injection) reveal a significant reduction in the number of activated macrophages. Green = F4/80; Grey = DAPI. Scale bar = 50μm Y-axis = % F4/80+ signal per total DAPI + signal in connective tissue area (*unpaired Student's t-test p<0.05). **(B)** μCT analysis of bone volume changes over time show PBS-lipo treated animals have a normal regeneration response which includes first a loss of bone volume followed by bone regeneration (n=4 mice, 16 digits). In contrast, Clo-Lipo treated animals exhibit a complete inhibition of both bone degradation and bone regrowth over the course of 35 days post amputation (n=4 mice, 16 digits) **(C)** 3D renderings of μCT scans enable visualization of the bone loss and regeneration in PBS-lipo treated animals. In Clo-lipo treated animals, there are no significant changes to bone architecture. For all images: distal = left, dorsal = top

To evaluate the effect of macrophage and osteoclast depletion on digit regeneration, we tracked anatomical and volumetric changes of the amputated P3 bone with µCT in Clo-Lipo and PBS-Lipo treated digits (Fig. 3B). 3D renderings of μCT scans show no change in bone architecture or bone volume over a 35 day period following amputation and Clo-Lipo treatment (Fig. 3B,C). Control digits exhibited a regeneration response similar to previous studies that included an initial bone volume decline prior to blastema formation followed by an average regrowth of 150% of the amputated stump bone volume (Fernando et al., 2011; Sammarco et al., 2015; Sammarco et al., 2014; Simkin et al., 2015b), thus indicating that the liposome vehicle did not alter the regenerative response (Fig. 3B,C). The data show that Clo-Lipo treatment inhibits both the bone degradation response and also the regeneration of distal bone. We also carried out studies to explore both the dose and timing of Clo-Lipo treatment of the regeneration response. A single injection of Clo-Lipo at the time of amputation resulted in digits that either failed to regenerate (37.5%; 3/8) or regenerated abnormally (62.5%; 5/8) producing boney spikes from regions of the stump (Suppl. Fig. 2A). Bone architecture suggests that the degradation phase is completely inhibited by a single treatment with Clo-Lipo, and that bone redifferentiation by the blastema is not an all or none event. To explore the relationship between the Clo-Lipo effect and the timing of the inflammation response, a single treatment with Clo-Lipo was administered at the peak of the inflammation response (7 DPA). Treatment at this time shows a trend toward reduced bone degradation and re-differentiation responses, but treated samples were not statistically different from PBS-Lipo treated controls (Suppl. Fig. 3, Two-way ANOVA main effect time, F=25.72, p<0.0001, and main effect treatment F=0.002, p=0.97). These studies identify the early stages of the inflammation response as being critical for the regeneration promoting effect that phagocytic cells, such as macrophages, have on blastema formation and digit tip regeneration in mice.

### Epidermal closure and histolysis is suspended by Clo-Lipo treatment

To understand better Clo-Lipo inhibited regeneration we carried out a histological analysis on samples that were treated with Clo-Lipo or PBS-Lipo and focused on key stages of the normal regenerative response (Fig. 4). PBS-Lipo treated digits demonstrate a regenerative response similar to untreated digits characterized in previous publications (Fernando et al., 2011; Sammarco et al., 2015; Sammarco et al., 2014; Simkin et al., 2015b). Briefly, at 7 DPA control digits show an open amputation wound, extensive stump bone degradation associated with osteoclast filled bone pits, a hypercellular bone marrow and activated osteoblast lining the endosteum and periosteum (Fig. 4A). At 10 DPA control digits have a closed wound epidermis, a prominent blastema and histological evidence of new osteoid deposition in the interface between the proximal blastema and the stump (Fig. 4B). At DPA 13 new bone growth in the form of woven bone islands are prominent at the blastema/stump interface (Fig. 4C arrowheads). At 28 DPA the regeneration of control digits is largely complete with the replacement of the digit tip that consist of newly regenerated woven bone, a reestablished distal marrow cavity, regenerated dermis and a regenerated distal nail plate (Fig. 4D).

**Figure 4.**
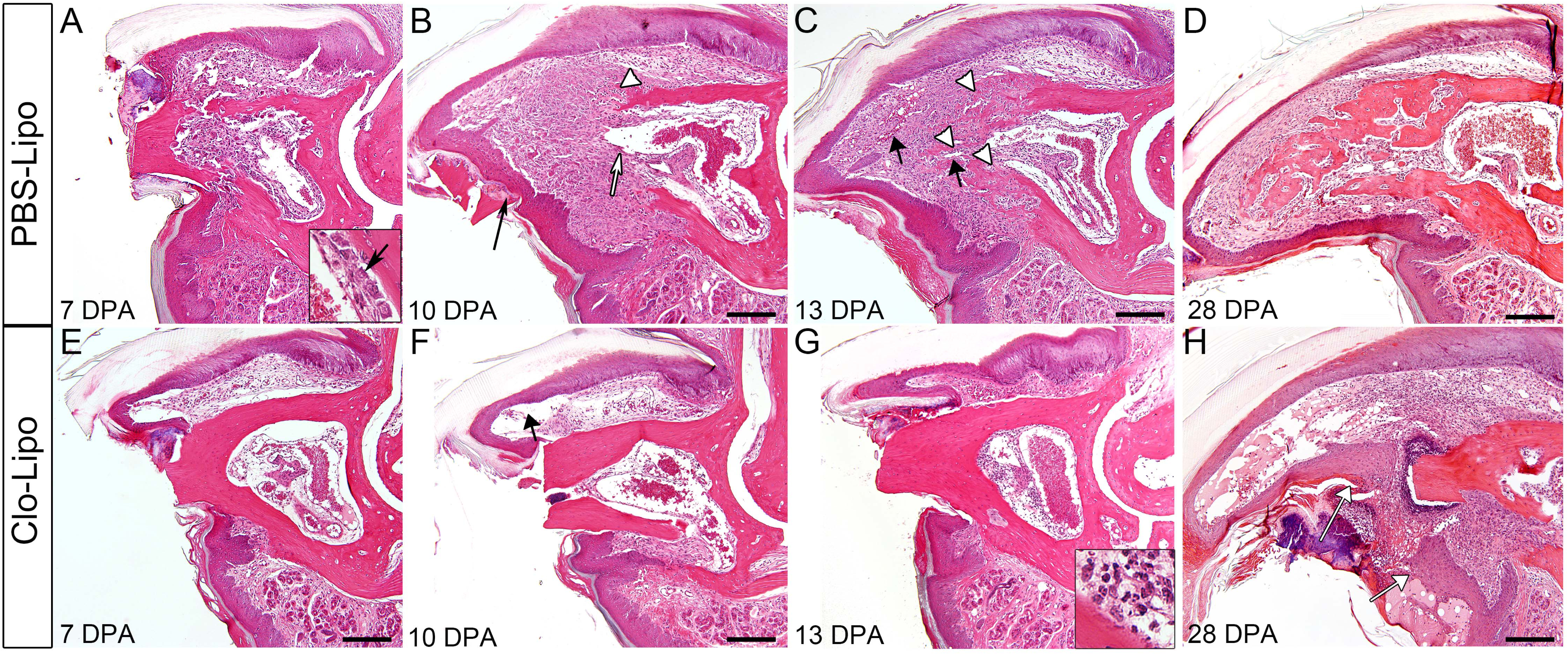
Injections of Clodronate liposomes inhibit histolysis and epidermal closure. **(A-D)** H&E staining of P3 digits over the course of regeneration in PBS-lipo treated digits. By 7 DPA (A), digits display bone degradation and endosteal activation (inset arrow). By 10 DPA (B) digits have a closed epidermis which ejects degraded bone fragments (black arrow), blastema formation contiguous with the bone marrow (white arrow) and the beginnings of new bone formation (white arrow head). By 13 DPA (C), new blood vessels have formed (black arrows) distal to the newly regenerating bone (white arrowheads). By 28 DPA (D), bone continues to be remodeled in trabecular islands. **(E-H)** H&E staining of P3 digits following Clo-Lipo injections show an open epidermis and lack of bone degradation at 7 DPA (E). Distal nail and epidermal growth (black arrow) is evident at 10 DPA (F) but not epidermal closure. At 13 DPA, there is a lack of re-epithelization, no bone degradation is observed and there is no blastema formation evident. Cell accumulation in the marrow consists of polymorphonuclear cells (inset). The epidermis has failed to close by 28 DPA (H white arrows) though distal epidermal and nail growth is evident. Scale bar = 200μm. For all images: distal = left, dorsal = top

In sharp contrast, the histology of Clo-Lipo treated digits show that the digit stump is largely unchanged during this 28 day period, and that the injury is suspended at an early phase of regeneration (Fig. 4E-H). There are a number of remarkable observations. First, there is no evidence of osteoclasts or bone pitting of the stump at any of the time points analyzed, and this is consistent with bone volume measurements from μCT imaging (Fig. 2B). Thus, Clo-Lipo treatment completely inhibits osteoclastogenesis and bone degradation typically associated with the regenerative response. Second, although nail elongation is observed, the epidermis fails to close over the amputation wound even by 28 DPA (Fig. 4H white arrows) indicating that Clo-Lipo treatment completely inhibits wound closure. By 7 DPA the epidermis is attached to the lateral periosteal bone surface as it does in controls (Fig. 4E), and this association with the periosteum is maintained while a longitudinal expansion of the epidermis creates a large pocket devoid of cells subjacent to the elongating nail (Fig. 4F, arrow). Thus, epidermal expansion and nail elongation does not appear to be dependent on macrophages. Third, we observe polymorphonuclear neutrophils associated with the amputation wound site (Fig. 4G inset) at all stages analyzed, suggesting that inflammation characteristics of the amputation wound is maintained and not completely resolved. Fourth, there is no histological evidence of blastema formation, indicating that Clo-Lipo treatment completely inhibits the distal accumulation of a regeneration blastema. Overall, these histological studies identify two obvious ways in which the depletion of macrophages could effectively inhibit the regenerative response: the absence of wound closure and/or the absence of osteoclast-mediated bone degradation.

### The specific loss of osteoclasts delay bone degradation and results in impaired regeneration

Osteoclasts are multinucleated cells of the monocyte lineage so their absence in Clo-Lipo studies was anticipated. To explore the hypothesis that osteoclasts, independent of macrophages, are required for regeneration we treated digits with free clodronate (F-Clo) to directly target osteoclasts (Russell and Rogers, 1999). To establish the efficacy of F-Clo, we administered a single injection of F-Clo into the digit at the time of amputation and evaluated osteoclast and macrophage presence at 7 DPA. Immunohistochemical studies show a distinct reduction of Cathepsin K positive osteoclasts when compared to PBS injected control digits (Fig. 5A,B). In contrast, F4/80 positive macrophages were abundant at the amputation wound site of F-Clo treated and PBS treated control digits (Fig. 5C,D). These data show that F-Clo treatment is effective in selectively depleting osteoclasts without impairing the macrophage population during digit tip regeneration, and this result is consistent with previous reports (Frith et al., 1997; Zeisberger et al., 2006). To study the effect of F-Clo on digit regeneration we measured changes in bone volume using μCT imaging following a single treatment with F-Clo at the time of amputation. Control digits injected with PBS undergo a normal regeneration response as previously described (see Fig. 2C,D). In contrast, digits receiving a single injection with F-Clo displayed an impaired regeneration response characterized by a delay in the onset of bone degradation and a reduced osteogenic response (Fig 5E,F; *Bonferroni post hoc test p<0.05, main effect treatment). Histological analysis at 14 DPA showed that wound closure is complete, and a blastema forms suggesting that the macrophage control of re-epithelization and blastema formation are independent of osteoclasts (Fig. 5G).

To address the necessity of bone degradation for regeneration we took advantage of our previous finding that the bone degradation phase is inhibited by wound closure and that application of the cyanoacrylic wound dressing Dermabond stimulates rapid re-epithelization (Simkin et al., 2015b). Based on histological analyses applying Dermabond to F-Clo-or PBS-treated digits stimulated epidermal closure by 7 DPA, and in both cases a blastema forms distal to the stump bone (Fig. 5H,I). MicroCT imaging of F-Clo/ Dermabond and PBS/ Dermabond treated digits showed that the amount of bone degradation was reduced in PBS digits and completely inhibited in F-Clo digits (Fig. 5J). Nevertheless, both PBS control and F-Clo treated digits regenerated the amputated digit tip indicating that digit tip regeneration occurs in the absence of stump bone degradation. Thus, the data indicate that while osteoclast-mediated bone degradation is a macrophage-dependent process, it is not a requirement for successful regeneration.

**Figure 5.**
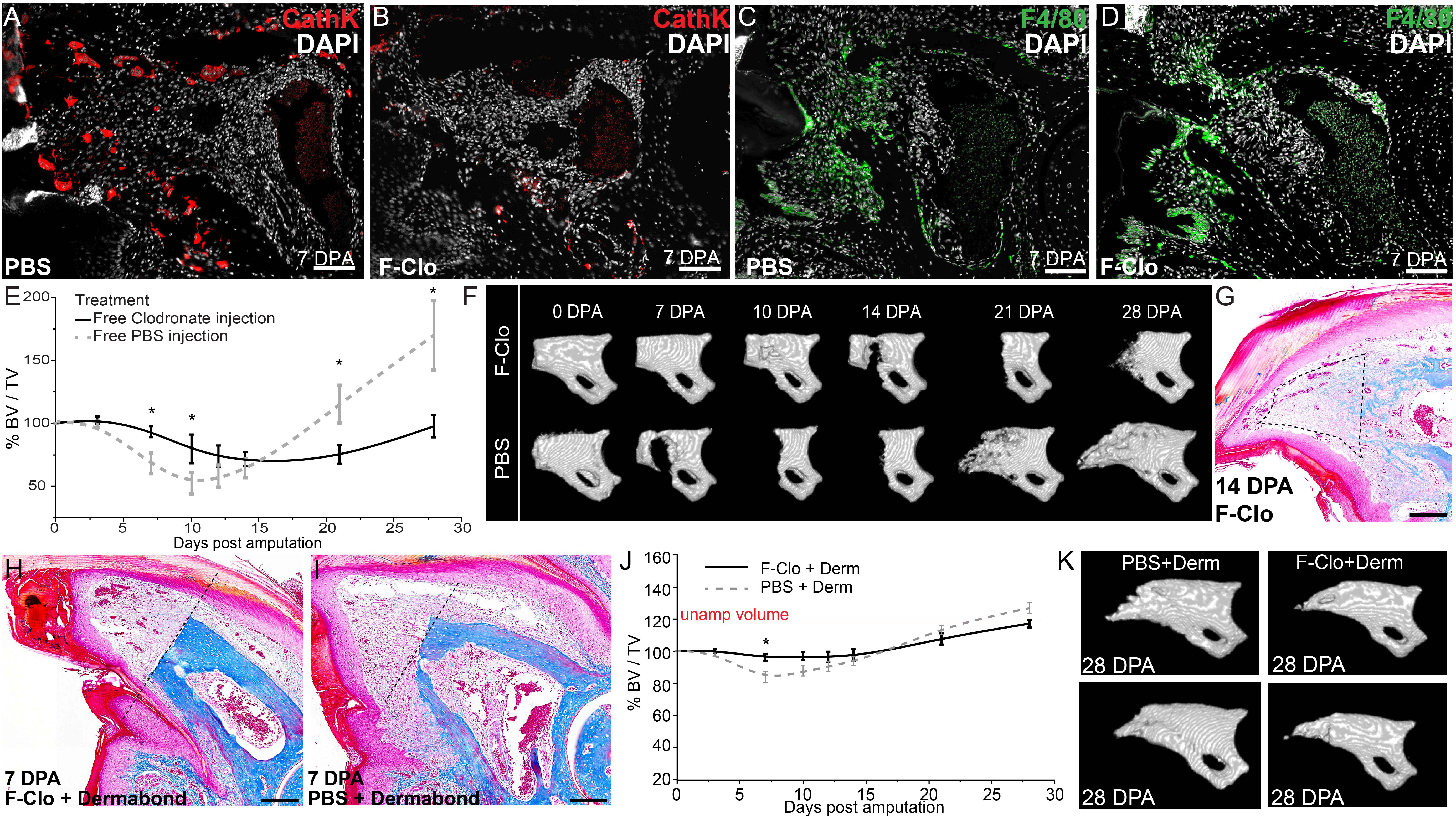
Osteoclast-specific depletion delays but does not inhibit regeneration. **(A-B)** A single injection of free clodronate (F-Clo) immediately following amputation depletes the P3 digit of Cathepsin K+ (CathK, red) cells. Representative image of a PBS-injected control digit (A) and a F-Clo injected digit (B) at 7 days post amputation (DPA). Scale bar = 100μm **(C-D)** F4/80+ staining (green) in PBS-injected digit (C) and F-Clo injected digit (D) at 7 DPA. Scale bar = 100μm. Grey = DAPI nuclear stain. **(E)** μCT analysis of bone volume changes with time after amputation in PBS treated digits (grey dotted line) and F-Clo treated digits (black line). Changes are measured in percent bone volume/total volume at time of amputation (%BV/TV). n=4 mice, 16 digits for both groups, *Bonferroni post hoc test p<0.05 main effect treatment at time points indicated. **(F)** 3D renderings of μCT images for PBS treated and F-Clo treated digits over time. **(G)** Trichrome staining of F-Clo treated digit at 14 DPA showing complete re-epithelization and accumulation of cells in the distal mesenchyme (dotted outline). Scale bar = 100μm. **(H-I)** Digits were treated with either F-Clo and Dermabond (H) or PBS and Dermabond (I). Trichrome staining reveals a loss of bone degradation from the original plane of amputation (black dotted line) in F-Clo+Derm treated digits and only minor degradation in PBS+Derm treated digits. Scale bar = 100μm **(J)** Micro-CT to track bone volume changes measured in percent bone volume/total volume at time of amputation (%BV/TV). Digits were treated with combined F-Clo and Dermabond (black line) or combined PBS and Dermabond (grey dotted line). In both treatment groups, bone regenerates to pre-amputation levels (red line). n=4 mice, 16 digits for both groups. *Bonferroni post hoc test p<0.05 main effect treatment. **(K)** Representative μCT images of P3 bone in either combined PBS and Dermabond treatment or combine F-Clo and Dermabond treatment groups at 28 DPA showing patterned bone growth in both groups. For all images: distal = left, dorsal = top

### Stimulating epidermal closure does not rescue Clo-lipo inhibition of regeneration

Because the wound epidermis is central to epimorphic regeneration in amphibian and zebrafish models, we next tested if re-epithelization could rescue regeneration in a macrophage-depleted, Clo-Lipo injected digit. We first established that Dermabond rescues the wound closure deficit that results from macrophage depletion and that re-epithelization is complete by 7 DPA (Fig. 6A,B). Clo-Lipo/Dermabond treated digits do not show evidence of bone degradation and develop a distal accumulation of cells that appear blastema-like (Fig. 6B). Next we used μCT imaging to track anatomical and bone volume changes of the digits during the regenerative response. In control studies combining Dermabond wound dressing with PBS-Lipo treatment we observed a regenerative response similar to that observed following only Dermabond treatment, whereas digits treated with Dermabond and Clo-Lipo displayed no change in stump bone volume or anatomy (Fig. 6C, D). These studies indicate that rescuing wound closure is not sufficient to rescue the regeneration response caused by macrophage depletion. The data show that the wound epidermis is able to recruit cells to the amputation wound to form a blastema-like structure, however maintenance of the blastema and the final stages of blastema differentiation is macrophage dependent. Overall, these studies indicate that macrophages are required for osteoclast-driven bone degradation, re-epithelization of the amputation wound and osteogenic differentiation of the blastema.

**Figure 6.**
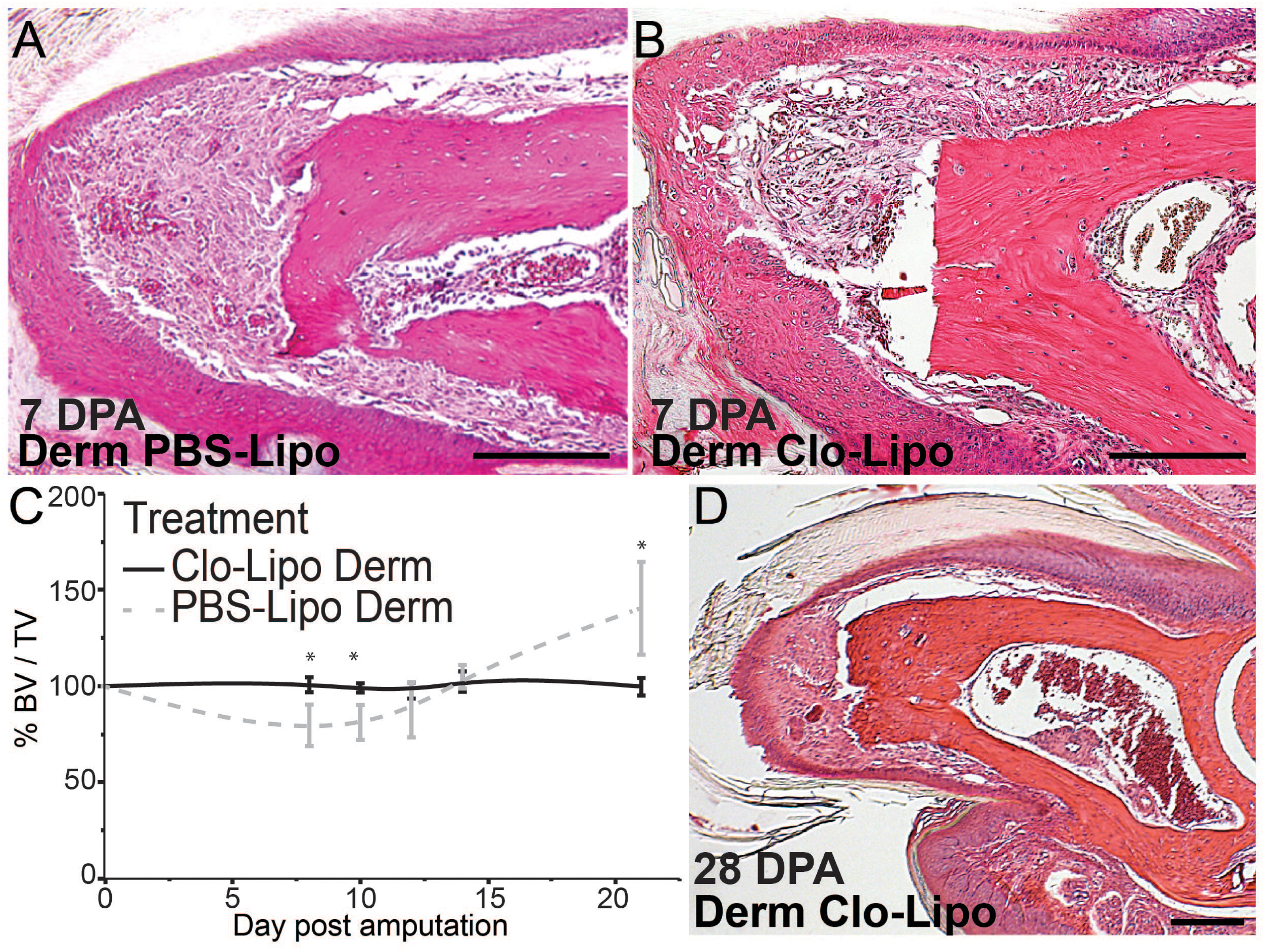
Rescuing epidermal closure does not rescue regeneration in Clo-Lipo treated digits. **(A–B)** H&E staining for histology shows application of Dermabond promotes epidermal closure and accumulation of mesenchymal cells under the epidermis by 7 DPA in PBS-Lipo treated digits (A) and Clo-Lipo treated digits (B). Scale bars = 100μm. **(C)** Micro-CT analysis of bone volume change over time reveals a loss of regeneration response in Clo-Lipo treated digits despite the rescue of epidermal closure. Y-axis = percent bone volume over total volume at time of amputation (%BV/TV), X-axis = days post amputation, n=8 digits for Clo-Lipo+Derm, n=4 digits for PBS-Lipo+Derm, (*Bonferroni post hoc test p<0.05, main effect treatment at timepoints indicated). **(D)** H&E of Dermbond treated, Clo-Lipo injected digits at 28 DPA shows no evidence of bone degradation or regrowth. Scale bar = 100μm. For all images: distal = left, dorsal = top

## DISCUSSION

In adult mammals, the resolution of traumatic injury throughout the body is tissue specific: some tissues undergo impaired healing with little sign of regeneration (e.g. skin, heart) whereas other tissues are able to regenerate a functional replacement (e.g. bone, skeletal muscle, liver) (Stocum, 2012). In all of these tissues the injury response involves inflammation that includes the mobilization and invasion of macrophages. The phagocytic action of macrophages functions to protect the host from infection although in recent years it is clear that macrophages also play key roles in enhancing the quality of the healing response and are necessary to promote regenerative responses. A key goal in regeneration biology is to understand the balance between the prevention of a pathological outcome resulting from uncontrolled infection or fibrosis on the one hand and enhancing the restoration of tissue function on the other. Both functions are critical for survival following traumatic injury, so both components are expected to be integral to the evolution of the healing response. Recent studies focusing on the inflammation response in animals that are highly regenerative demonstrate that macrophage recruitment to an amputation wound is required for a successful regenerative response (Godwin et al., 2013; Petrie et al., 2014). In the current study we have used the regenerating digit tip of the mouse to show that macrophage invasion of the amputation wound in mice is absolutely essential for multiple stages of regeneration. Macrophage depletion following digit amputation causes numerous modifications of the regeneration program that include an inhibition in osteoclastogenesis and the bone degradation phase of regeneration, an inhibition of re-epithelization of the amputation wound, the inhibition of blastema formation, and the inhibition of redifferentiation of the blastema to reform the digit tip. Our studies show that while macrophages are required for osteoclastogenesis and bone degradation, this process itself is not required for a successful regenerative response. On the other hand, macrophages are essential for re-epithelization of the wound and the wound epidermis is required for blastema formation and regeneration. Finally, macrophages are required for blastema cell differentiation to regenerate distal skeletal elements. Overall, the evidence shows that epimorphic regeneration in a mammalian model is macrophage-dependent and that macrophages regulate multiple key components of the regeneration response.

### Osteoclasts and bone degradation are not required for regeneration

Blastema formation during digit tip regeneration is characterized by the dramatic histolytic degradation of the P3 bone stump that occurs prior to wound closure. This response is associated with an increase in osteoclast numbers that coincides with the peak of invading macrophages and results in a significant reduction in the volume of the stump bone (Fernando et al., 2011). Following the local depletion of macrophages, we find few osteoclasts associated with the stump bone and a complete absence of the degradation response. Thus, the rise in stump osteoclasts and the subsequent degradation of the P3 stump bone are dependent on local macrophages during the early stages following amputation injury. These findings are in line with similar macrophage depletion studies that result in a loss of osteoclasts in bone regeneration models associated with fracture healing (Lin et al., 2016). Osteoclasts are large multinucleated cells derived from the monocyte/macrophage lineage and form from the fusion of single nucleated pre-osteoclasts. Osteoclasts are generally viewed as a bone-specific resident monocyte population (Sinder et al., 2015), however we find few F4/80 or CD45 positive cells associated with the P3 skeletal element prior to amputation. The paucity of resident cells coupled with the dramatic rise in macrophages is consistent with the conclusion that the majority of macrophages are recruited to the amputation wound. Since osteoclasts associated with bone degradation following amputation are lacking after local macrophage depletion with Clo-Lipo treatment, we conclude that these multinucleated cells are derived, at least in part, from a recruited macrophage population.

While it is clear that osteoclasts play an obvious histolytic role in the regeneration response, it has not been clear whether the degradation of bone is required for a regenerative response. There is evidence implicating the bone degradation response in regulating blastema size and the extent of the regenerative response (Sammarco et al., 2015). Studies in other models of epimorphic regeneration show that histolytic activity is up-regulated early in regeneration and that matrix metalloproteinase activity plays a role in regenerate patterning (Bai et al., 2005; Grillo et al., 1968; Vinarsky et al., 2005; Yang and Bryant, 1994; Yang et al., 1999). In mouse digit regeneration we have shown that wound closure signals the termination of the bone degradation phase and transitions the regeneration response to blastema formation (Simkin et al., 2015b). Here we show that the use of clodronate to selectively deplete osteoclasts and delay degradation combined with enhancing wound closure to precociously terminate degradation completely eliminates bone degradation without depleting the macrophage population. The results show clearly that blastema formation and regeneration still occurs under conditions in which there is no anatomical evidence of bone degradation, thus it is clear that histolysis of the bone stump, although a prominent feature of the regeneration response, is not required for an epimorphic regenerative response in mice. In parallel, fracture healing studies show that osteoclast specific depletion in a tissue-specific model of bone regeneration is not required for the osteogenic response (Alexander et al., 2011).

### The wound epidermis is required for blastema accumulation

Local depletion of macrophages during digit tip regeneration causes a complete inhibition of re-epithelization of the amputation wound. Similar results from tissue-specific wound healing studies have been reported (Leibovich and Ross, 1975; Lucas et al., 2010; Mirza et al., 2009) indicating that macrophages are required for wound closure. One difference between digit amputation and full thickness wounds is that re-epithelization of full thickness wounds is delayed but eventually occurs and this likely reflects differences in wound contraction between the two models, i.e. healing of digit amputations does not involve wound contraction (Fernando et al., 2011). In digit amputation, we have previously shown that re-epithelization can be enhanced by simply treating the amputation wound with a cyanoacrylic wound dressing, Dermabond, (Simkin et al., 2015b) and we show in the current study that Dermabond completely rescues re-epithelization inhibited by macrophage depletion. Such rescue is consistent with the conclusion that macrophages play a role in creating a wound environment permissive for re-epithelization rather than having a direct effect on epidermal cells. The lack of an apparent effect on epidermal expansion and nail elongation in macrophage-depleted digits also supports this conclusion. Oxygen availability and epidermal hypoxia have been implicated in the regulation of wound epidermis formation (Simkin 2015; Sammarco 2015). Oxygen availability is generally mediated by the vasculature and following traumatic injury oxygen availability is largely regulated by the angiogenic response. In wound healing models macrophages infiltrate the granulation tissue of the wound bed where they create a proangiogenic environment that results in an excessive revasularization response that has been linked to scar formation (DiPietro, 2016). Angiogenesis during digit blastema formation is uniquely different in that the injury response is not characterized by a proangiogenic environment, but an anti-angiogenic environment that is regulated by the production of the anti-angiogenic factor, PEDF, coupled with a reduced level of *Vegfa* expression (Muneoka et al., 2008; Yu et al., 2014). In this light, the observation that the digit blastema is largely devoid of macrophages is consistent with both the avascular and hypoxic character of the blastema (Fernando et al., 2011; Sammarco et al., 2014).

The necessity of the wound epidermal and mesenchymal cell interactions for epimorphic regeneration is established in other models such as salamander limb regeneration and zebrafish fin regeneration (Carlson, 1969; Chablais and Jazwinska, 2010; Goss, 1956; Mescher, 1976; Whitehead et al., 2005). In mammalian regeneration there is little direct evidence that the wound epidermis plays a similar role. The observation that macrophage depletion inhibits the formation of the wound epidermis, blastema formation and blastema differentiation has allowed us to explore the relationship between these events. Stimulating the formation of the wound epidermis following osteoclast-depletion rescues the regenerative response but does not rescue regeneration in macrophage-depleted digits. This provides evidence that the mammalian wound epidermis is required for a successful regeneration response. Even though regeneration is not rescued by the wound epidermis in macrophage-depleted digits, the wound epidermis does stimulate the accumulation of a population of cells that appear blastema-like distal to the amputation stump. This indicates that the wound epidermis plays a role in recruiting cells that accumulate distal to the amputation injury, although establishing the regenerative potential of this structure will require further examination. The wound epidermis along with the deviation of a nerve plays a similar role in salamanders since wounding and nerve deviation stimulates the formation of an ectopic blastemal structure that eventually regresses (Endo et al., 2004; Satoh et al., 2008). Based on these findings we conclude that the necessity of the wound epidermis for epimorphic regeneration can be extended to mammals, and represents an example of a regenerative mechanism that is conserved among vertebrates.

### Macrophages play multiple roles in epimorphic regeneration

It is difficult to gain a clear understanding of the role that macrophages play in epimorphic regeneration based solely on macrophage depletion studies. Nevertheless, a major conclusion is that macrophages are required for multiple key aspects of the regeneration response in mammals, and this study provides an important first step toward elucidating the details and extent of their involvement. In addition to a role in regulating osteoclast-driven bone histolysis and a role in stimulating wound re-epithelization, macrophages are required for the final stages of blastema differentiation to reform the digit tip. Since digit tip regeneration involved direct ossification of blastema cells to form woven bone, we conclude that macrophages are required for bone regeneration. This conclusion is consistent with fracture healing studies where macrophages are interlaced throughout regions where woven bone forms by direct ossification, and bone regeneration is inhibited following macrophage depletion (Alexander et al., 2011). Curiously, the spatial-temporal profile of macrophages during digit tip regeneration does not correlate with regions of skeletal differentiation of the blastema. Macrophages numbers peak early in wound healing prior to histological evidence of blastema formation, and macrophages are largely absent in the blastema where osteogenesis is occurring. Thus, in digit tip regeneration the macrophage effect on skeletal differentiation must occur early in the osteogenic process. The localization of macrophages proximal to the blastema and prior to blastema formation raises the possibility that they play a role in modulating the recruitment of osteoprogenitor cells from the stump into the blastema. Further studies are required to clarify the specific role that macrophages play in osteogenesis during blastema differentiation.

During traumatic injury the inflammatory response must navigate a fine balance between the initial protection against infection versus the eventual promotion of a functional repair response (Godwin et al., 2016). In classical epimorphic regeneration models, such as the salamander limb or the zebrafish fin, recent macrophage depletion studies provide clear evidence that this balance is tipped toward the promotion of a functional regeneration response (Godwin et al., 2013; Petrie et al., 2014), and our study adds the mouse digit tip to this list of macrophage-dependent regenerative responses. Epimorphic regeneration in adult mammals is relatively rare whereas the regeneration of specific tissues such as muscle and bone can be quite robust. Other tissues, such as the skin, display regenerative responses only during specific developmental stages (fetal) or in selective regions of the body (e.g. oral skin), whereas adult skin typically undergoes a non-regenerative healing response that culminates in the deposition of scar tissue (Mak et al., 2009; Martin and Leibovich, 2005). It is interesting that the inflammation response is known to promote the regeneration of bone and muscle tissue, while inhibiting regeneration during skin healing (Martin et al., 2003; Mori et al., 2008; Novak et al., 2014; Raggatt et al., 2014). These observations point to the evolution of an inflammatory balance between the initial protection against infection that drive the response in skin wounds versus the promotion of regenerative responses in internal tissues like muscle and bone. It is interesting that some of the macrophage activities identified in epimorphic regeneration parallel established tissue-specific responses of skin (e.g. promotion of re-epithelization) and bone (e.g. promotion of osteogenesis). Thus the data supports the idea that what makes epimorphic regeneration unique is the way in which multiple tissue-specific responses are coordinated both temporally and spatially, and that macrophages play a key role in this process. This conclusion also helps to bridge the interface between epimorphic and tissue-specific regenerative responses, and provides an avenue for the development of strategies to enhance regeneration in mammals.

## MATERIALS AND METHODS

### Digit amputations and animal care

Adult 8-week old female CD1 mice were obtained from Charles River Laboratories (Wilmington, MA). Mice were anesthetized with 1-5% isofluorane gas with continuous inhalation. The second and fourth digits of both hind limbs were amputated at the P3 distal level as described previously (Simkin et al., 2013). Digits were collected at specified time points for histological analysis. All experiments were performed in accordance with the standard operating procedures approved by the Institutional Animal Care and Use Committees at Tulane University Health Sciences Center and the College of Veterinary Medicine at Texas A&M University.

### Histology and Immunohistochemistry

Tissue was harvested at specified time points and fixed in Zinc buffered formalin (Anatech, Battle Creek, MI) overnight. Bone was decalcified for 8 hours in formic acid based decalcifier (Decal I, Surgipath, Richmond, IL). Samples were processed for paraffin embedding using a Leica TP 1020 Processor (Leica, Buffalo Grove, IL). 4 μm serial sections were obtained using a Leica RM2255 microtome. Sections were deparaffinized in xylenes and rehydrated through a series of graded ethanol. Mayer's Hematoxylin and Eosin Y (Sigma-Aldrich, St. Louis, MO) staining was carried out according to manufacturer's protocol. Mallory Trichrome staining was also carried out according to manufacturer's instructions (American Mastertech, Lodi, CA). Coverslips were mounted with Permount mounting medium (Fisher Scientific, Waltham, MA). For immunohistochemistry, serial sections were deparaffinized in xylene and rehydrated through graded ethanol. Antigen retrieval was carried out in either a pH6 citrate buffer for 20 minutes at 90°C or with proteinase K at 10mg/mL for 10 minutes at 37°C according to in-house optimized protocols for each antibody. Endogenous hydrogen peroxide was blocked using a solution of 3% H_2_O_2_ in methanol, and endogenous avidin and biotin were blocked with a Dako blocking kit. Non-specific antibody binding sites were blocked using a serum free blocking buffer (Dako, Carpinteria, CA). Slides were incubated at 4°C overnight with the following primary antibodies: F4/80 (5μg/mL, Rat anti mouse, Cat# 14-4801, eBioscience, San Diego, CA), Ly6B.2 (0.1μg/mL, Rat anti mouse, Cat# MCA771A, AbD Serotec/BioRad), Cathepsin K (2μg/mL, Rabbit anti mouse, Cat# 19027, Abcam, Cambridge, MA), CD45 (5μg/mL, Rat anti mouse, Cat# 103101, BioLegend, San Diego, CA). Primary antibody detection was carried out using either secondary antibodies conjugated to Alexa Fluor 488 and 568 (Invitrogen, Carlsbad, CA) or secondary antibodies conjugated to biotin and resolved with a tyramide amplification kit according to manufacturer's instructions (TSA kit #T20912, Invitrogen / ThermoFisher Scientific).

### Image analysis

Brightfield images of histological sections were obtained using a 10x, 20x or 40x objective on an Olympus BX60 upright microscope equipped with an Olympus DP72 camera. Fluorescent micrographs were acquired on an Olympus BX61 fluorescence deconvolution microscope. Quantification of fluorescent signal was performed using masking subsampling of positive fluorescent area in Slidebook Imaging Software (Intelligent Imagine Innovations, Denver, CO). Total area of 488 or 568nm fluorescent signal was calculated and normalized to total DAPI area to calculate % positive area / total cellular area. Signal quantification was restricted to the connective tissue area, excluding nail, epidermis, scab, bone and bone marrow. The entire P3 area of a representative section of each sample was imaged for quantification at 10x magnification. Autofluorescent red blood cells were subtracted from images using a Slidebook subtraction algorithm before quantification to reduce background signal.

### MCP-1 bead implants

400μm Cibacron blue Affi-gel agarose beads (BioRad) are soaked in 0.5 mg/mL MCP1 (Prospec, Cat# CHM-313) with 0.1% BSA in PBS. Vehicle control beads were soaked in 0.1% BSA in PBS. Bead implants were carried out as previously described (Simkin et al., 2013). Briefly, after soaking in protein solution overnight, beads were allowed to air dry and were implanted with tungsten needles into the dermis surrounding the P3 bone at 0 or 3 DPA. For each treated mouse, digits on one paw received PBS soaked beads and on the other paw MCP1 soaked beads. Left/right paw treatment was randomized for each mouse. Macrophage recruitment was calculated with immunofluorescent analysis as described above.

### Osteoclast, Macrophage depletion and rescue of re-epithelization

For macrophage or osteoclast depletion, 10μL of 50mg/mL Clodronate liposomes (Clo-Lipo), or PBS liposomes (PBS-Lipo, www.ClodronateLiposomes.com) was injected into the P2 region of each amputated digit using an insulin syringe. Each digit received an injection at 0, 2, and 5 DPA. For depletion of osteoclasts, Clodronate (1.85μg/g body weight) was injected in 10μL of PBS into the P2 region just prior to digit amputation. For control digits, 10μL of PBS was injected alone. Depletion efficacy was quantified by immunofluorescence for Cathepsin K and F4/80+ cells. For rescue of wound closure experiments, 10μL of Dermabond (Ethicon, LLC, San Lorenzo, Puerto Rico) was applied to each digit immediately following amputation. Digits were allowed to dry for 1 minute following Dermabond application and then injected with Clo-Lipo, PBS-lipo, F-Clo or PBS.

### Micro-CT analysis

Micro-CT images were acquired using a VivaCT 40 (Scanco Medical AG, Bruttisellen, Switzerland) at 1000 projections per 180 degrees with a voxel resolution of 10μm^3^, and energy and intensity settings of 55V and 145μA respectively. Integration time for capturing the projections was set to 380 ms using continuous rotation. Images were segmented using the BoneJ (Doube et al., 2010) (Version 1.2.1) Optimize Threshold Plugin for ImageJ (Version1.48c). Changes in bone volume were quantified using the BoneJ Volume Fraction Plugin for ImageJ. Percent bone volume divided by total bone volume (%BV/TV) was calculated by normalizing the measurement of each digit to its original volume immediately following amputation. Final images were compiled using Adobe Photoshop CS4 and CS6.

### Statistical analysis

Bone volume graphs were compiled and were analyzed using Two-way ANOVA with main effects treatment and time using JMP (v. 10.0.0, SAS Institute Inc.). Bonferroni multiple comparison tests were conducted for simple effect treatment at specific time points when appropriate and reported on the graphs. Graphs of immuno-positive area for cell counting studies were compiled and analyzed using Prism (version 6, Graphpad). Two-way ANOVA with main effects time and treatment or unpaired Student's t-test for simple effect treatment were calculated as indicated in figure legends. All figures were compiled using Adobe Photoshop and Adobe Illustrator (Creative Suite 6).

## ACKNOWLEDGEMENTS

The authors thank members of the Muneoka lab, Larry Suva and Dana Gaddy for discussions. Funding for this project was provided by W911NF-06-1-0161 from DARPA, W911NF-09-1-0305 from the US Army Research Laboratory, the John L. and Mary Wright Ebaugh endowment fund at Tulane University, and Texas A&M University. The authors have no conflicts of interest to declare.

## SUPPLEMENTAL FIGURES

**Supplemental Figure 1. Clo-Lipo injections do not deplete osteoblast or neutrophil populations. (A)** Map of amputated digit at 5 DPA showing area represented in images (B) and (C). **(B)** 1 day post Clo-Lipo injection, Osx+ cells are still present and line the periosteum of the amputated bone, however the Osx+ cells maintain a squamous morphology associated with quiescent osteoblasts. **(C)** Immunohistochemical stain for the neutrophil cell surface marker Ly6B.2. Neutrophils are still present at the injury site 1 day after the final Clo-Lipo injections (6 days post amputation).

**Supplemental Figure 2. A single injection of Clo-Lipo or PBS-Lipo at 0 DPA immediately following amputation results in partial inhibition of regeneration. (A)** 3D renderings of μCT scansof Clo-Lipo treated digits at 21 days post amputation (DPA). 3 out of 8 digits treated with Clo-Lipo at 0 DPA show no degradation or new bone growth by 21 DPA whereas 5/8 digits show unpatterned bone growth. **(B)** Digits treated with PBS-Lipo at 0 DPA show patterned bone growth by 21 DPA.

**Supplemental Figure 3**. A single injection of Clo-Lipo at 7 DPA does not inhibit regeneration. Clo-Lipo or PBS-Lipo was injected at 7 DPA (arrow) when macrophages and osteoclasts are at peak activity. Clo-Lipo treated digits (black line) show a trend toward less bone growth compared to PBS-Lipo treated digits (grey dotted line) but differences in final volumes are not statistically significant (Two-way ANOVA main effect time, F=25.72, p<0.0001, and main effect treatment F=0.002, p=0.97). Y-axis = %bone volume / total volume at time of amputation. X-axis = Days post amputation

